## supplemental file for "Non-glycosylated IGF2 prohormones are more mitogenic than native IGF2"

### Contents

**Figure S1.** RP-HPLC chromatograms of purified IGF2 proforms 3

**Figure S2.** MS spectra of purified IGF2 proforms 4

Determination of secondary structures of IGF2 and IGF2 prohormones by Circular Dichroism and theoretical calculations 6

**Table S1.** Secondary structure content (in %) 8

**Figure S3.** Comparison of IGF2 NMR structure with predicted structures of IGF2 and big-IGF2(87) 9

**Figure S4.** Analysis of predicted IGF2 structure 9

**Figure S5.** Analysis of structures predicted using AlphaFold2 10

**Table S2.** Secondary structure content (in %) 10

**Table S3.** Receptor-binding affinities 11

**Figure S6.** Binding curves for (A) IR-A, (B) IR-B, (C) IGF1-R, (D) R- 12

**Figure S7.** Saturation binding curve of [$^{125}$I]-monoiodotyrosyl-Tyr2-IGF2 and [$^{125}$I]-monoiodotyrosyl-TyrA14-insulin on R-cells 13

**Figure S8.** Saturation binding curve of [$^{125}$I]-monoiodotyrosyl-Tyr2-IGF2 on immobilized IGFBP3 13

**Figure S9.** A typical saturation binding curve of [$^{125}$I]-monoiodotyrosyl-Tyr2-IGF2 on immobilized IGFBP3:ALS complex 14

**Figure S10.** Binding curves of ligands on (A) D11:IGF2R, (B) IGFBP3, (C) ALS:IGFBP3 15

**Figure S11.** Representative Western blots showing expression of particular receptors in different cells. 16

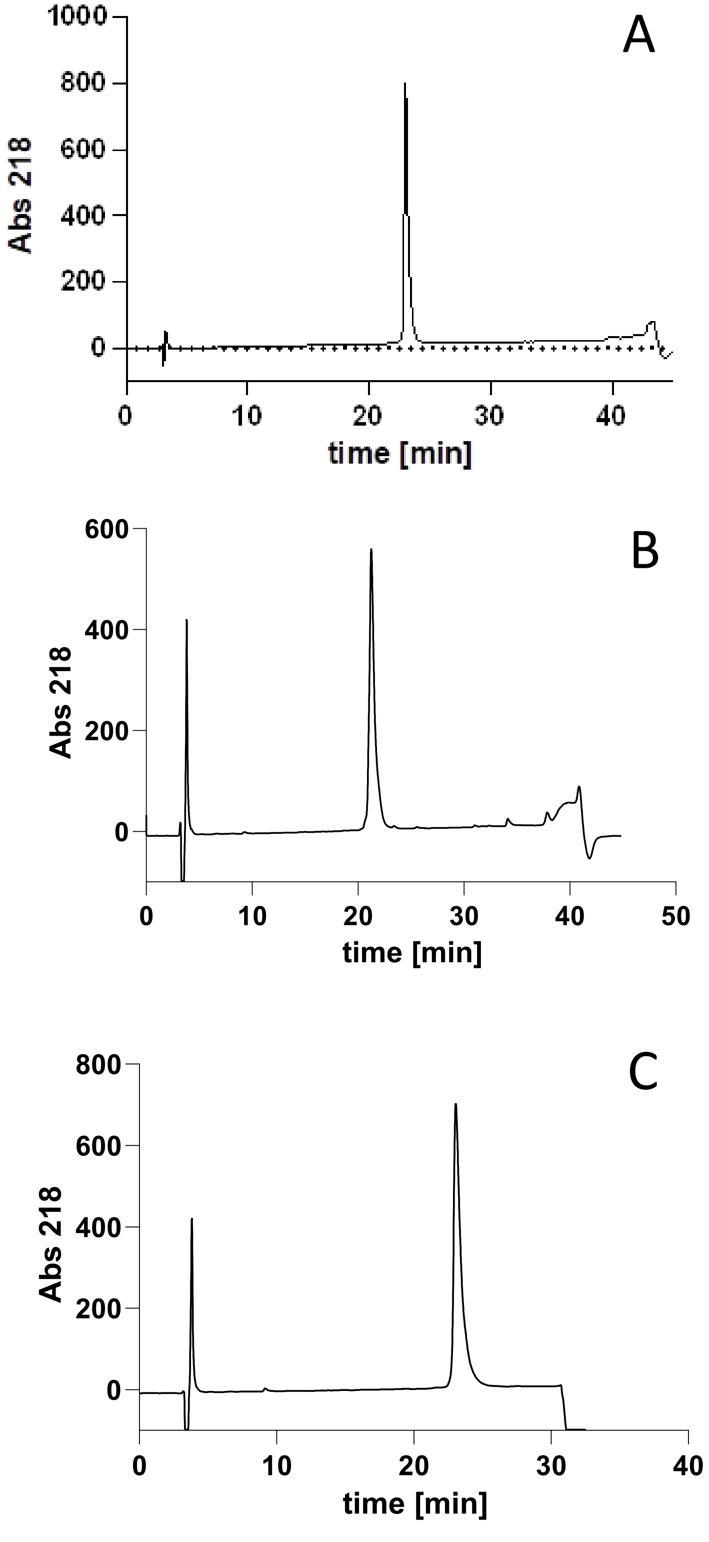


**Figure S1.** RP-HPLC chromatogram of the purified (**A**) big-IGF2(87), (**B**) big-IGF2(104), and (**C**) pro-IGF2(156). Analysis was done on Vydac C4 column 0.4 x 25 cm in a gradient of acetonitrile in water with 0.1 % TFA.

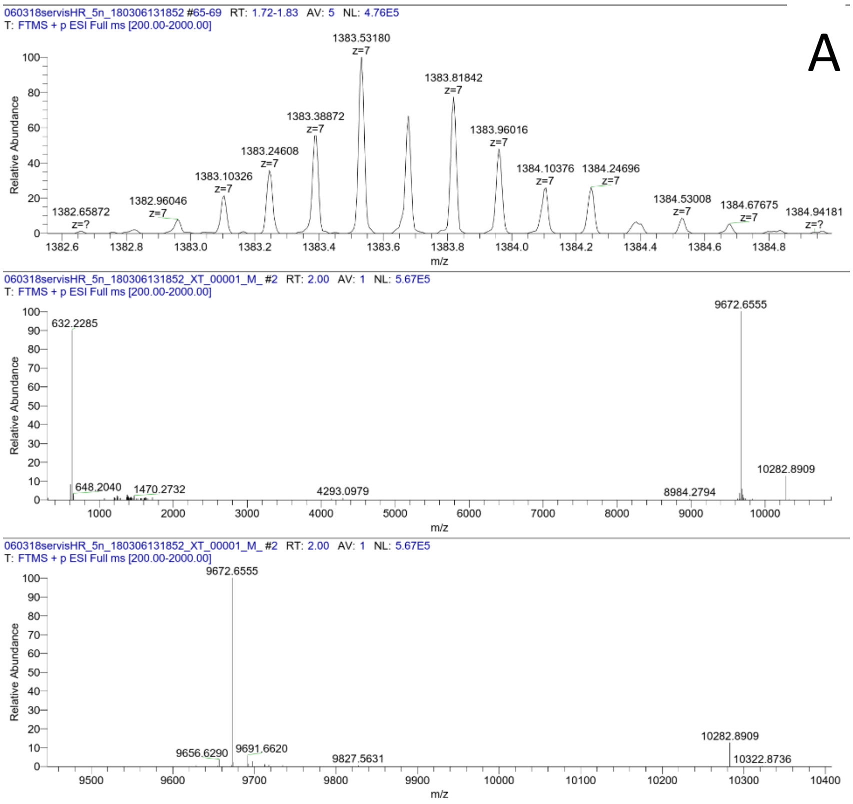
A
060318servisHR_5n_180306131852 #65-69 RT: 1.72-1.83 AV: 5 NL: 4.76E5
T: FTMS + p ESI Full ms [200.00-2000.00]
Relative Abundance
1382.65872 z=?
1382.96046 z=7
1383.10326 z=7
1383.24608 z=7
1383.38872 z=7
1383.53180 z=7
1383.81842 z=7
1383.96016 z=7
1384.10376 z=7
1384.24696 z=7
1384.53008 z=7
1384.67675 z=7
1384.94181 z=?
m/z
060318servisHR_5n_180306131852_XT_00001_M_ #2 RT: 2.00 AV: 1 NL: 5.67E5
T: FTMS + p ESI Full ms [200.00-2000.00]
632.2285
648.2040
1470.2732
4293.0979
8984.2794
9672.6555
10282.8909
9656.6290
9691.6620
9827.5631
10322.8736

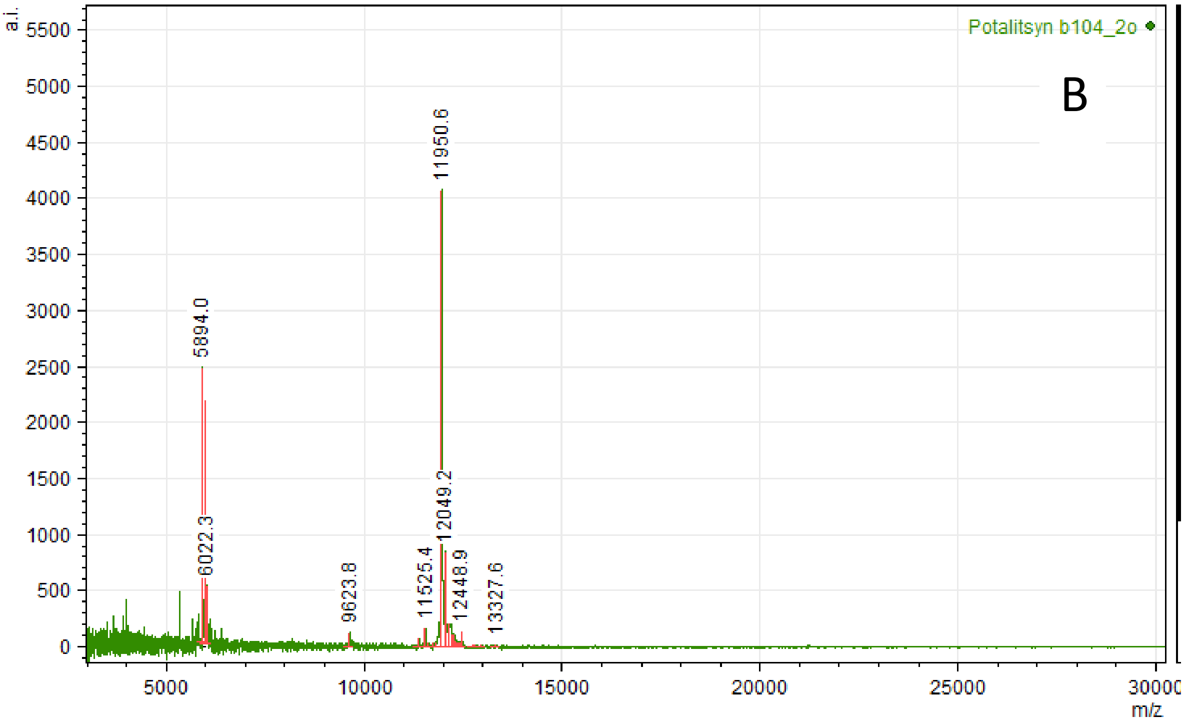
B
Potalitsyn b104_2o
a.i.
5894.0
6022.3
9623.8
11525.4
11950.6
12049.2
12448.9
13327.6
m/z

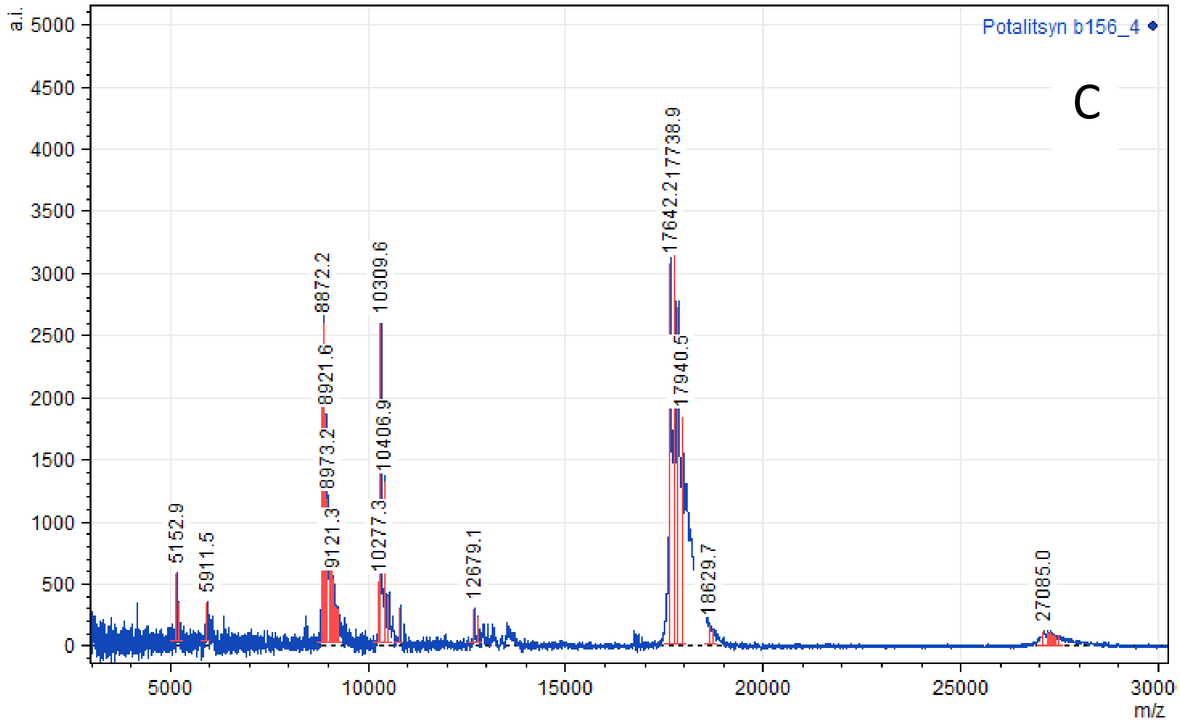


**Figure S2.** (**A**) ESI mass spectrum of big-IGF2(87). Expected molecular weight is 9672, (**B**) MALDI mass spectrum of big-IGF2(104). Expected molecular weight is 11955.5, (**C**) MALDI mass spectrum of pro-IGF2(156). Expected molecular weight is 17640.2.

***Determination of secondary structures of IGF2 and IGF2 prohormones by Circular Dichroism and theoretical calculations***

All spectra were normalized to concentration, optical path length, and the number of amino acids in a particular protein. For IGF2, we can observe a +/-/- pattern with maximum at ~191 nm and minima at ~207 and 223 nm. This corresponds to the presence of a segment with helical conformation in the protein. Nevertheless, the intensity is substantially lower than is typical for highly helical proteins (Greenfield, 2006). Therefore, the portion of the α-helical part within the whole IGF2 can be assumed as ~20 %. Big-IGF2(87) and big-IGF2(104) exhibit similar CD patterns to IGF2, only the intensity is slightly lower. Therefore, their α-helical content will obviously be lower. The largest protein in our study, pro-IGF2(156), exhibited a significantly distinct spectrum with zero signal (or only weakly positive) at 185-190 nm and two negative minima at ~203 and ~225 nm. This would suggest another reduction of the α-helical content in the protein and an increase in the contribution of other secondary structures.

Secondary structure content (SSC) was more accurately estimated by decomposing individual CD spectra, using the BeStSel (Micsonai, 2022; (Micsonai, 2018) and K2D3 programs (Kabsch, 1983). We observed (BeStSel) a gradual decrease of the helical content with the increasing size of the protein, from 25 % for IGF2 to 13 % for pro-IGF2(156). This is partially compensated for by the increasing content of β-strand or "other" conformations. SSC estimated using K2D3 differs from BeStSel. However, K2D3 restricts the analysis to the 190-240 nm region, whereas BeStSel uses a slightly extended region, and therefore we consider the BeStSel estimates to be more reliable. For a detailed analysis, see text and Table S2.

The observed decrease of the helical content for bigger proteins is simply due to an increase in their size with an unchanged proportion of helical content. This means that only parts between Ala1 and Glu67 hold some secondary structure, while the rest is probably unstructured. This was confirmed by calculating the minimal helical content $hc$ (see Methods and Table S3 for details) that corresponds relatively well to α-helix SSC obtained with the BeStSel analysis. We also tried to predict 3D structures of big-IGF2s and pro-IGF2(156) (and IGF2 as a control) using ColabFold (Mirdita, 2022), which combines the homology search of MMseq2 (Mirdita, 2019) with AlphaFold2 (Jumper, 2021). Figure S3A shows an overlay of the predicted IGF2 structure with the reported NMR structure (2L29), revealing their high degree of resemblance that is confirmed by the analysis of the structures (Figure S4). Calculated $hc$ for predicted IGF2 corresponds well to that estimated by CD (or to the minimal $hc$; see Table S3). Analysis of the predicted structure of big-IGF2(87) (cf. Figures S4B and S5) and its calculated $hc$ of 26 % suggest that it also corresponds well to the actual protein structure, even though the accuracy rate decreases starting from a position around ~60. Only the parts of big-IGF2(104) and pro-IGF2(156) corresponding to IGF2 (Ala1 - Glu67) were reliably predicted (Figures S5A-F), including the relative positions of the helical segments (Figures S5G-I). Models predicted additional helical segments for big-IGF2(104) (Phe90-Leu102) and pro-IGF2(156) (Tyr92-Arg130), resulting in unrealisticaly high $hc$ of 33 % and 38 %, respectively. To conclude, the secondary structure between Ala1 and Glu67 in big-IGF2s and pro-IGF2 are largely similar to IGF2, whereas their remaining parts are rather unstructured.

To interpret the CD spectra of our proteins in greater depth, we estimated the secondary structure content (SSC), using the BeStSel (Micsonai, 2022) and K2D3 (Louis-Jeune, 2012) programs. Both programs approximate the experimental CD spectra by a linear combination of a set of basic spectra, corresponding to proteins with a solved structure and thus with a known secondary structure composition. BeStSel uses 8 secondary structure motifs (2 a-helices, 3 antiparallel and 1 parallel b-strands, 1 turn, and others; according to (KabschSander, 1983). The K2D3 program only distinguishes a-helical, b-stranded, and “other” conformations. Note that neither method is able to recognize other helical conformations (e.g. PPII, $3_{10}$) and these are gathered as “others”. Table S1 summarizes the estimated SSC for all proteins and both methods. We can see a gradual decrease of the helical content with increasing size of the protein. This is compensated for by the increasing content of b-strand or others (depending on the method). As K2D3 limits the analysis to the region of 190-240 nm, while BeStSel uses an extended region of 185-250 nm, we find the BeStSel estimations more reliable. For brevity, we have grouped both a-helices into one family of conformation, as well as all b-strands. Figure 1 presents these grouped SSC estimates.

The observed decrease of the helical content for bigger proteins is simply due to an increase in their size, with an unchanged proportion of helical content. This means that only parts between ALA1 and GLU67 hold some secondary structure, while the rest is probably unstructured (or contains secondary structures labeled as “others”). We calculated the minimal helical content $hc$ (see Table S2), based precisely on the assumption that the proportion of the helix is unchanged, and only the size of the protein increases. We obtained $hc$ of 27-36 % (IGF2), 21-28 % (big-IGF2(87)), 17-23 % (big-IGF2(104)), and 12-15 % (pro-IGF2(156)). Specific $hc$ (and its range) depends on the reference structure used. Obtained $hc$ corresponds relatively well to a-helix SSC obtained with BeStSel, which supports the idea of an unstructured "tail" of big-IGFs.

**Table S1.** Secondary structure content (in %) estimated for all studied proteins according to experimental CD.

| | | | **BeStSel** | | |
|---|---|---|---|---|---|
| **185-250 nm** | | **IGF2** | **big-IGF2(87)** | **big-IGF2(104)** | **pro-IGF2(156)** |
| α-helix | regular | 14 | 12 | 11 | 5 |
| | distorted | 11 | 11 | 10 | 8 |
| | sum | 25 | 23 | 21 | 13 |
| β-strand | antiparallel | 18 | 18 | 18 | 23 |
| | parallel | 0 | 0 | 0 | 0 |
| | sum | 18 | 18 | 18 | 23 |
| turn | | 15 | 14 | 14 | 17 |
| others | | 42 | 45 | 47 | 48 |
| | | | **K2D3** | | |
| **190-240 nm** | | | | | |
| α-helix | | 18 | 14 | 12 | 3 |
| β-strand | | 21 | 21 | 24 | 32 |
| others | | 61 | 65 | 64 | 65 |

#### ***Alpha fold predictions***

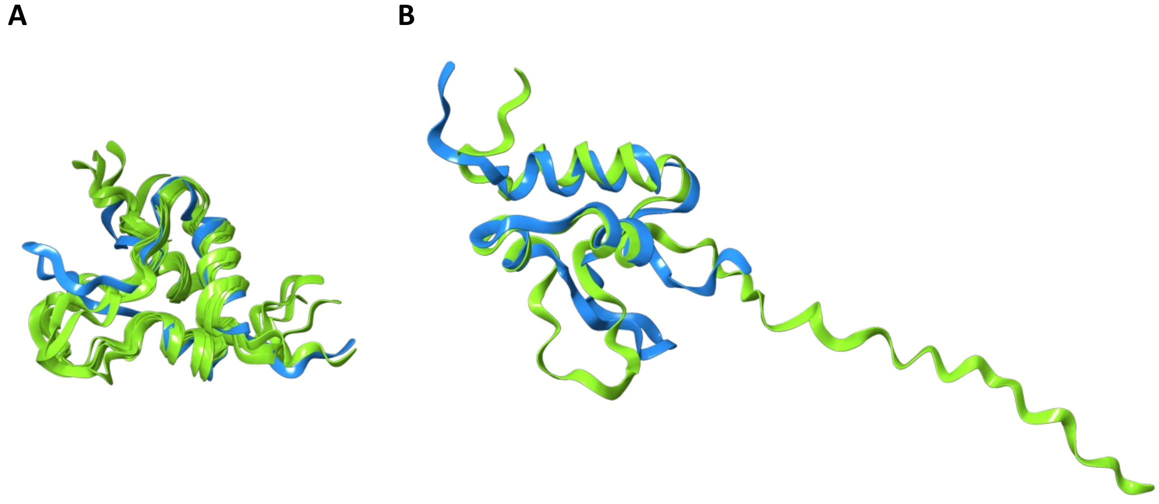


**Figure S3.** Comparison of IGF2 NMR structure (2L29; blue structures) with predicted (AlphaFold2; green) structure of (**A**) five predicted structures of IGF2 and (**B**) big-IGF2(87).

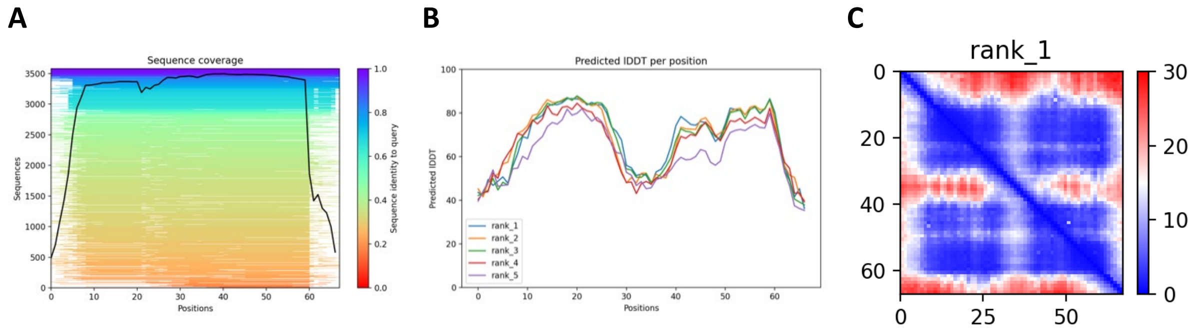


**Figure S4.** Analysis of predicted (AlphaFold2) IGF2 structure used as a control: the sequence coverage diagram (**A**), predicted LDDT curves for five IGF2 structures (**B**), predicted aligned error for the highest ranked IGF2 structure (**C**).

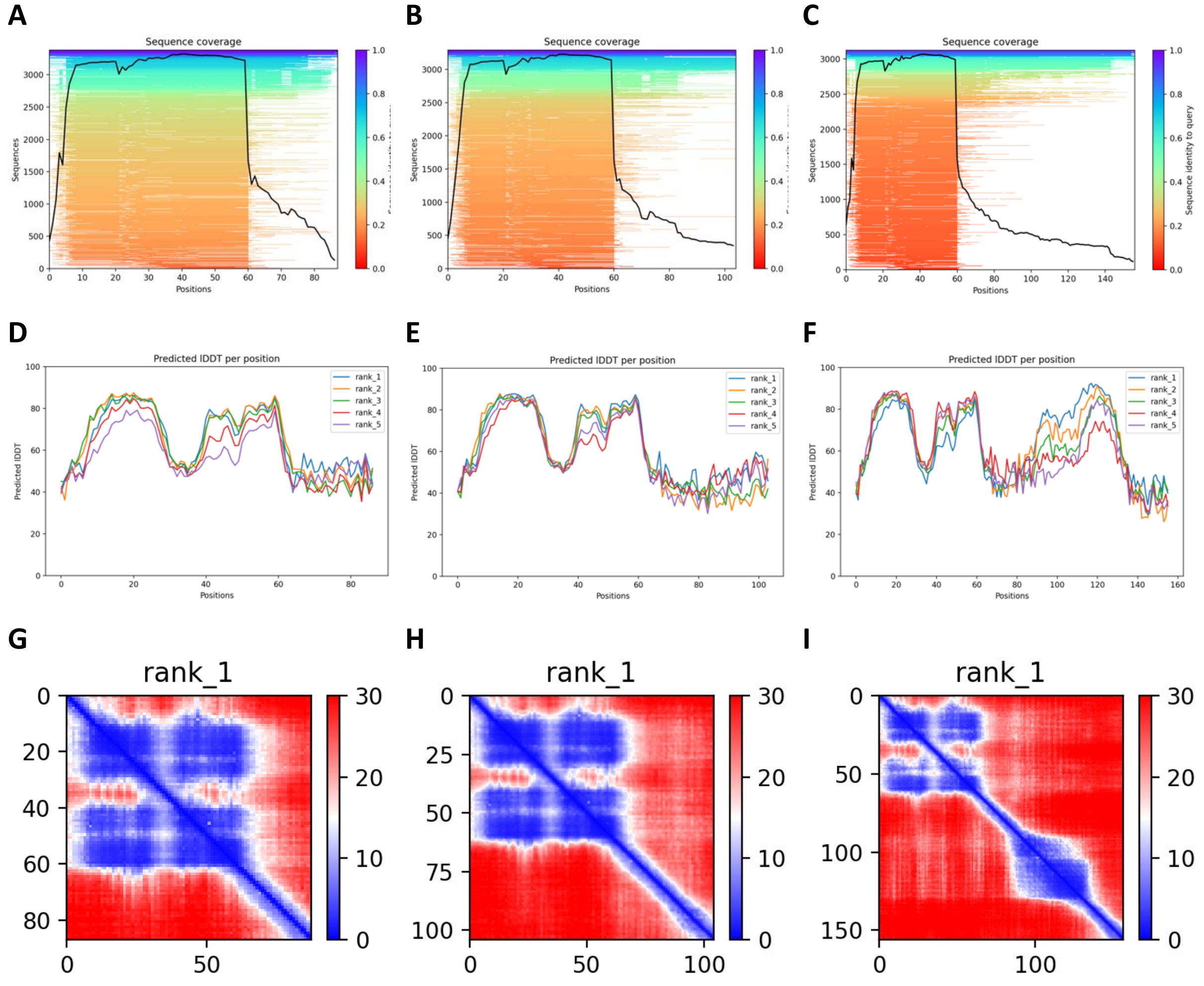


**Figure S5.** Analysis of structures predicted using AlphaFold2. Top row: the sequence coverage diagram of (**A**) big-IGF2(87), (**B**) big-IGF2(104), and (**C**) pro-IGF2(156); Middle row: predicted LDDT curves for five structures of (**D**) big-IGF2(87), (**E**) big-IGF2(104), and (**F**) pro-IGF2(156); Bottom row: predicted aligned error for the highest ranked structure of (**G**) big-IGF2(87), (**H**) big-IGF2(104), and (**I**) pro-IGF2(156).

**Table S2.** Secondary structure content (in %) estimated for all studied proteins using different methods

| | helical content (%) | | | |
|---|---|---|---|---|
| **Model** | **IGF2** | **big-IGF2(87)** | **big-IGF2(104)** | **pro-IGF2(156)** |
| CD (BeStSel) | 25 | 23 | 21 | 13 |
| Minimal *hc* | 27-36 | 21-28 | 17-23 | 12-15 |
| AlphaFold2 | 33 | 26 | 33 | 38 |

[a]*hc* = number of amino acids in $\alpha$-helical conformation as in IGF2 (pdb structures - 2L29, 2V5P, 6VWG) / number of all amino acids (67, 87, 104, and 156). The wider range of values is due to the unequal number of "helical" amino acids in different pdb structures.

**Table S3.** Receptor-binding affinities of IGF1, IGF2, human insulin, big-IGF2(87), big-IGF2(104), and pro-IGF2(156) on individual receptors or binding proteins. The $K_d$ values of individual proteins were determined in (n) independent series of measurements. The affinity of the native hormones for their cognate receptors was set at 100 % and relative binding affinities of ligands were calculated and are shown in the main text (Table 1).

| | IR-A<br>$K_d$ ± S.D. (*n*) | IR-B<br>$K_d$ ± S.D. (*n*) | IGF1R<br>$K_d$ ± S.D. (*n*) | IGFBP3<br>$K_d$ ± S.D. (*n*) | D11:IGF2R<br>$K_d$ ± S.D. (*n*) | M6P/IGF2R<br>$K_d$ ± S.D. (*n*) | IGFBP3:ALS<br>$K_d$ ± S.D. (*n*) |
|---|---|---|---|---|---|---|---|
| IGF2 | 3.17 ± 1.25 (3) (100%) | 7.02 ± 2.93 (3) (100%) | 0.76 ± 0.12[a] (3) | 0.2 ± 0.10 (4) (100%) | 1.29 ± 0.27 (7) (100%) | 0.94 ± 0.36 (4) (100%) | 0.95 ± 0.31 (3) (100%) |
| IGF1 | 23.8 ± 6.6 (3)* | 224 ± 16 (4)* | 0.12 ± 0.02[a] (3) (100%)<br>0.24 ± 0.08[b] (5) (100%) | 0.31 ± 0.07 (3) | n.b. | 31.4 ± 2.61 (3) | n.d. |
| HI | 0.27 ± 0.06 (4) | 0.26 ± 0.07 (4) | 292 ± 31[b] (3)* | n.d. | n.d. | n.d. | n.d. |
| Big-IGF2(87) | 3.82 ± 0.23 (3) | 9.47 ± 3.30 (3) | 1.60 ± 0.55[b] (5) | 0.29 ± 0.17 (3) | 1.77 ± 0.74 (4) | 0.56 ± 0.25 (5) | 1.63 ± 0.99 (3) |
| Big-IGF2(104) | 2.75 ± 0.28 (3) | 8.83 ± 2.74 (3) | 2.48 ± 0.32[b] (6) | 0.14 ± 0.05 (3) | 0.19 ± 0.07 (4) | 0.14 ± 0.03 (6) | 5.58 ± 0.99 (3) |
| Pro-IGF2(156) | 12.21 ± 1.93 (3) | 16.04 ± 3.97 (2) | 67.9 ± 27.2[b] (5) | 1.25 ± 0.63 (3) | 2.31 ± 1.68 (3) | 6.33 ± 4.71 (4) | 10.5 ± 3.14 (3) |

The $K_d$ values and were calculated from at least three independent measurements (n, number of replicates). [a, b] The individual $K_d$ values of ligands in this column were determined in two independent series of experiments and are relative to a corresponding native IGF1 $K_d$ value. ([a] to [a] and [b] to [b]). n.b. means no binding. n.d. means not determined. *Data from Ref. (Krizkova, 2016).

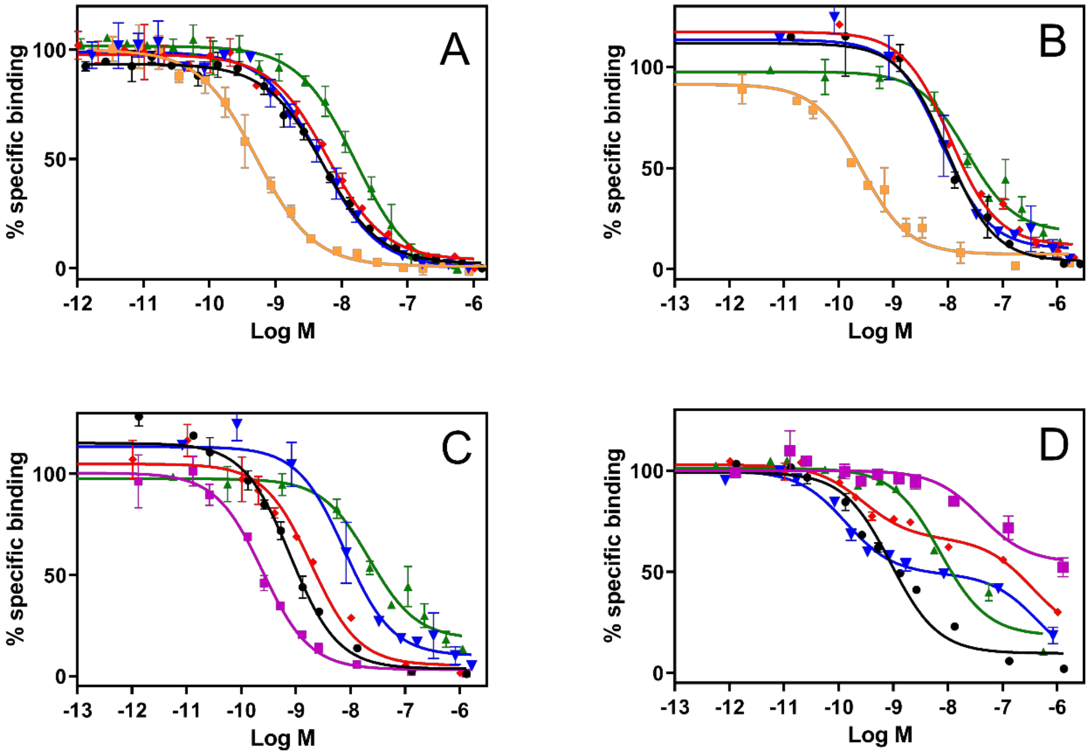


**Figure S6.** Binding curves for (**A**) IR-A, (**B**) IR-B, (**C**) IGF1-R, (**D**) R-. Inhibition of binding of human [$^{125}$I]-monoiodotyrosyl-ligand to corresponding receptor by human insulin (orange), IGF1 (magenta), IGF2 (black), big-IGF2(87) (red), big-IGF2(104) (blue), pro-IGF2(156) (green). Representative binding curve for each hormone or analog is shown.

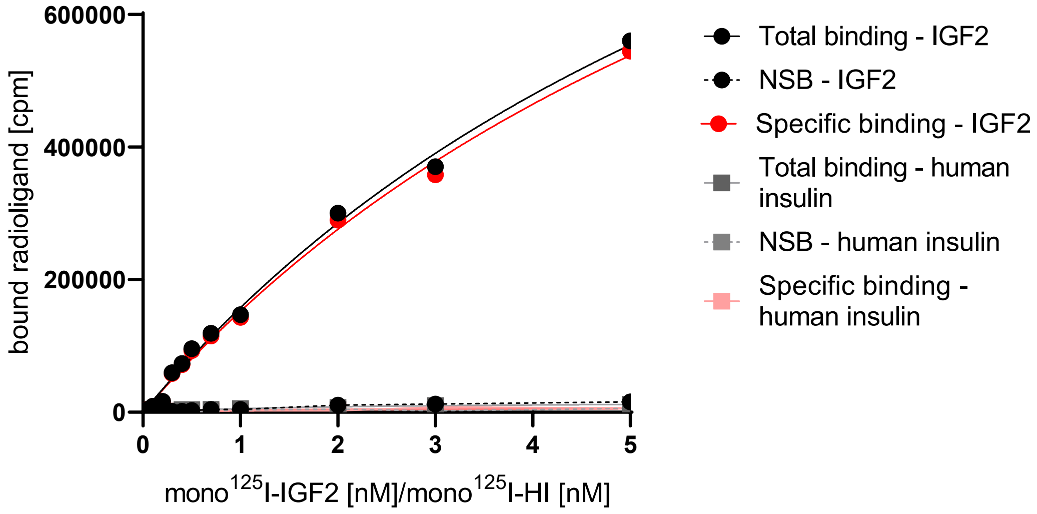


**Figure S7.** A typical saturation binding curve of [$^{125}$I]-monoiodotyrosyl-Tyr2-IGF2 and [$^{125}$I]-monoiodotyrosyl-TyrA14-HI to R-cells. Statistical analysis of the results from three such independent experiments provided the final $K_d$ value 8.20 ± 0.85 nM (n = 3) for IGF2 to IGF2R.

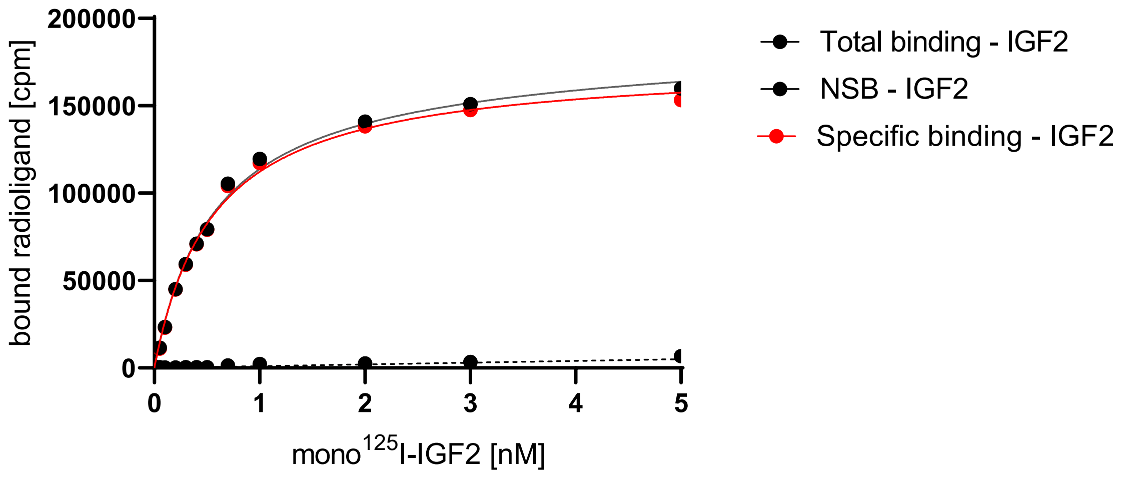


**Figure S8.** A typical saturation binding curve of [$^{125}$I]-monoiodotyrosyl-Tyr2-IGF2 to IGFBP3. Statistical analysis of the results from three such independent experiments provided the final $K_d$ value 0.59 ± 0.29 nM (n = 3) for IGF2 to IGFBP3.

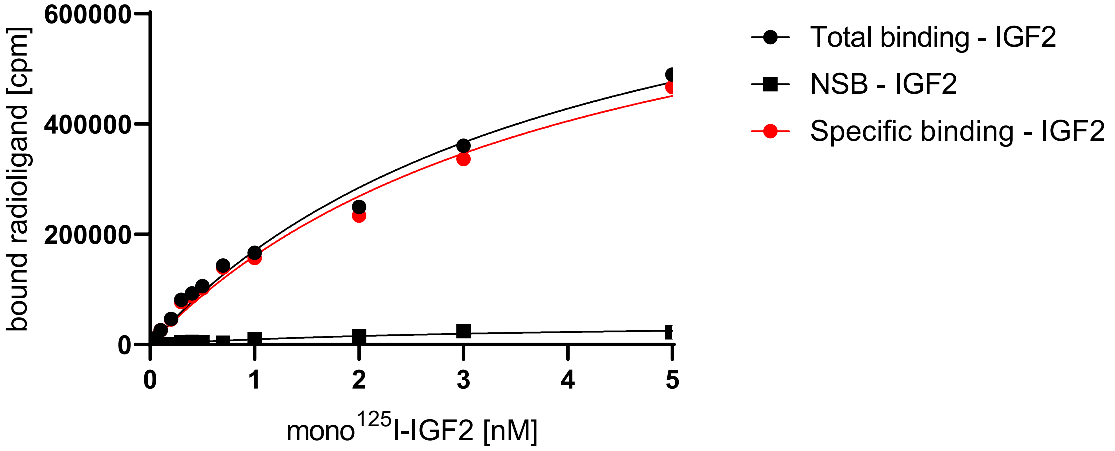


**Figure S9.** A typical saturation binding curve of [$^{125}$I]-monoiodotyrosyl-Tyr2-IGF2 with IGFBP3 to ALS. Two independent measurements gave us the mean $K_d$ value. $K_d$ value was set at value 5.90 ± 2.53 nM (n = 2) for IGF2:IGFPB3 to ALS.

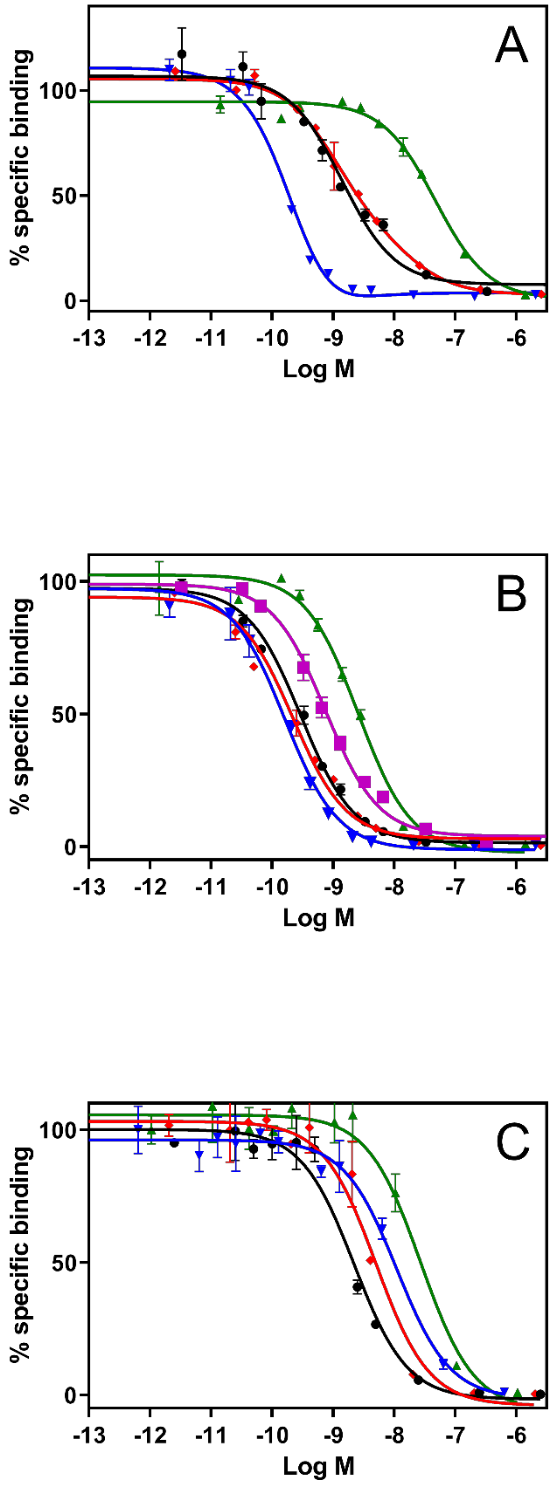


**Figure S10.** Binding curves for (**A**) D11:IGF2R, (**B**) IGFBP3, (**C**) ALS:IGFBP3. Inhibition of binding of human [$^{125}$I]-monoiodotyrosyl-ligand to corresponding receptor by IGF1 (magenta), IGF2 (black), big-IGF2(87) (red), big-IGF2(104) (blue), pro-IGF2(156) (green). Representative binding curve for each hormone or analog is shown.

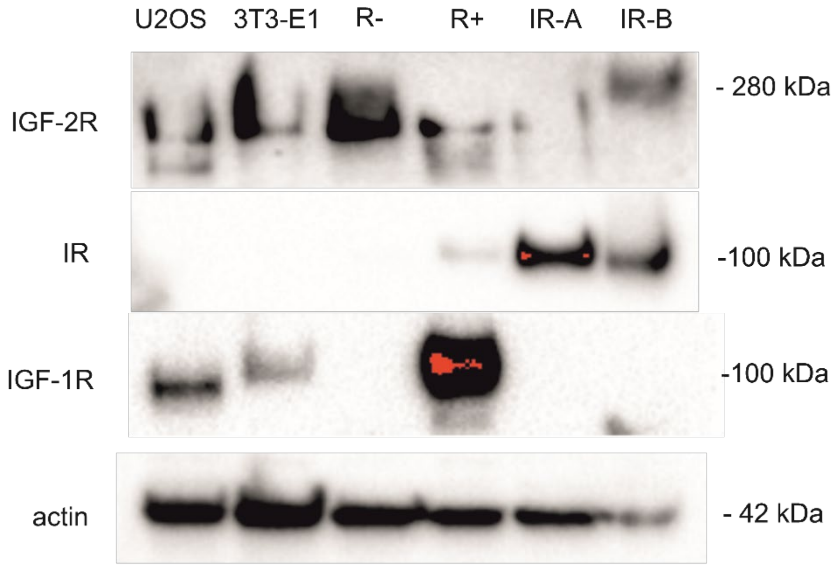


**Figure S11.** Representative Western blots showing expression of particular receptors in different cells.
